# Nf2 fine-tunes proliferation and tissue alignment during closure of the optic fissure in the embryonic mouse eye

**DOI:** 10.1101/2020.06.28.176065

**Authors:** Wesley R. Sun, Sara Ramirez, Kelly E. Spiller, Yan Zhao, Sabine Fuhrmann

## Abstract

Uveal coloboma represents one of the most common congenital ocular malformations accounting for up to 10% of childhood blindness (1~ in 5,000 live birth). Coloboma originates from defective fusion of the optic fissure (OF), a transient gap that forms during eye morphogenesis by asymmetric, ventral invagination. Genetic heterogeneity combined with the activity of developmentally regulated genes suggest multiple mechanisms regulating OF closure. The tumor suppressor and FERM domain protein neurofibromin 2 (NF2) controls diverse processes in cancer, development and regeneration, via Hippo pathway and cytoskeleton regulation. In humans, *NF2* mutations can cause ocular abnormalities, including coloboma, however, its actual role in OF closure is unknown. Using conditional inactivation in the embryonic mouse eye, our data indicates that loss of *Nf2* function results in a novel underlying cause for coloboma. In particular, mutant eyes show substantially increased RPE proliferation in the fissure region with concomitant acquisition of RPE cell fate. Cells lining the OF margin can maintain RPE fate ectopically and fail to transition from neuroepithelial to cuboidal shape. In the dorsal RPE of the optic cup, *Nf2* inactivation leads to a robust increase in cell number, with local disorganization of the cytoskeleton components F-actin and pMLC2. We propose that RPE hyperproliferation is the primary cause for the observed defects causing insufficient alignment of the OF margins in *Nf2* mutants and failure to fuse properly, resulting in persistent coloboma. Our findings indicate that limiting proliferation particularly in the RPE layer is a critical mechanism during optic fissure closure.

## Introduction

Congenital ocular malformations such as anophthalmia, microphthalmia and coloboma (MAC) are the cause for over 25% of childhood blindness worldwide (1–3). Uveal coloboma alone may account for up to 10% of childhood blindness, 1~ in 5,000 live birth (4, 5) for reviews, see (6–8); it is most commonly observed as missing tissue in the ventro-nasal region of the eye and originates from a failure in apposition and fusion of the margins of the optic fissure (OF). The OF forms as a ventral groove by asymmetric invagination, extending from the vitreal side to the proximal junction with the forebrain that allows mesenchymal cells to migrate in and form the hyaloid vessel (9–11). Closure of the OF occurs when mesenchymal cells end migration; the laterally growing edges of the retinal pigmented epithelium (RPE) and retina cells lining the OF margins appose to fuse and form a continuous optic cup. Fusion starts at the midpoint of the OF and moves in both directions distally and proximally (9, 12). The etiology of coloboma is complex, and defects result from disturbance of heterogenous genetic and/or environmental factors suggesting multiple mechanisms in regulating the closure process, including cell autonomous and cell non-autonomous tissue interactions. In humans, a subpopulation of colobomata is a result of mutations in developmentally important genes, however, the origins for many OF closure defects are unknown (for reviews, see (3, 7, 8, 13, 14)). Additional genes have been identified in animal models; substantial progress has been made in understanding critical processes, including growth and patterning of the ventral optic cup and optic nerve head (15–19), cell-cell contact and signaling (20–30), crosstalk with migrating hyaloid precursors and extracellular matrix components (31–36), cytoskeleton dynamics (37, 38), epigenetics (39), degradation of ECM and cellular proteins (40–42), programmed cell death, survival and cell proliferation (43, 44) (for reviews, see (7, 8, 13)). Elegant in vivo imaging studies in zebrafish and excellent anatomical analyses in chick have characterized important morphogenetic and cellular behavior (20, 27, 34, 45–49). In addition, comprehensive gene expression analyses have identified novel candidate genes potentially relevant for regulating OF closure (27, 47, 50–52).

Gene mutations of Hippo signaling pathway components represent novel associations with coloboma (7, 23, 53, 54). Mutations in the Neurofibromin 2 gene (*NF2*) result in the autosomal dominant disorder Neurofibromatosis Type 2 (NF2, OMIM # 101000), a relatively rare syndrome characteristic of neoplastic lesions. Human *NF2* mutations associate frequently with ocular abnormalities (53, 55–57). Importantly, some patients with loss of *NF2* heterozygosity also exhibit coloboma, consistent with *Nf2* being identified as a coloboma gene in mouse (53, 58, 59) (this study). The *Nf2* gene encodes the FERM domain protein MERLIN, affiliated with Ezrin, Radixin, and Moesin (ERM) proteins. It exerts multiple functions in tumor suppression, development, and regeneration in diverse organs and tissues, for example by restricting proliferation, controlling apoptosis, and promoting differentiation, apicobasal polarity and junctional complex formation (60–62). NF2 localizes primarily to the cell cortex and links the cortical actomyosin cytoskeleton to the plasma membrane. It associates with α-catenin to promote establishment of intercellular junctions and polarity complexes (63). In addition, NF2 can localize to the nucleus and act as a contact-dependent tumor suppressor by inhibiting CRL4DCAF1 ubiquitin ligase (61). Importantly, among other regulatory inputs, NF2 also controls the Hippo pathway to balance organ size, proliferation and behavior of stem cells; it binds to and activates several Hippo pathway components, including the Hippo kinases MST1/2 and LATS1/2 (61, 62). Loss of Hippo signaling leads to nuclear translocation of the transcriptional co-activators YAP/TAZ, often associated with abnormal proliferation in cancer and during development. Human mutations of *YAP* and its transcriptional effector *TEAD* can result in coloboma and the autosomal dominant ocular disorder Sveinsson’s chorioretinal atrophy, respectively (SCRA, OMIM # 108985) (54, 64, 65). Recent studies in Drosophila, zebrafish and mouse elucidate functions of Yap/Taz during ocular development, with some aspects of evolutionary conservation (59, 66–71) (for reviews, see (72–74)).

Our study focuses on the role of Nf2 during OF closure in mouse. NF2 protein is expressed in developing ocular tissues and is required for development of the lens and the optic cup, particularly of the ciliary margin into ciliary body and iris (58, 59, 75–77). Coloboma occurs upon tissue-specific *Nf2* deletion (58, 59), however, it is unclear how loss of *Nf2* function affects the closure process. Our data demonstrates that *Nf2* disruption results in ectopic proliferation in the RPE layer in the OF region with concomitant maintenance of RPE cell fate. RPE cells in the temporal OF margin maintain columnar organization instead of transitioning to cuboidal shape. In the dorsal RPE of the optic cup, *Nf2* loss of function leads to a substantial increase in cell number, accompanied with local disorganization of cytoskeleton components F-actin and pMLC2. We propose that these abnormalities in the *Nf2* mutant optic cup cause insufficient alignment of the OF margins, resulting in a failure to attach and fuse properly, and, thus, leading to a persistent coloboma. Our analysis provides novel insights into the underlying mechanisms during OF closure as well as into the specific function of Nf2 in this process.

## Results

### Loss of *Nf2* function leads to a narrow coloboma in the ventral optic cup

Germline disruption of the *Nf2* gene causes early embryonic lethality before eye development commences (78). Therefore, we conditionally disrupted *Nf2* using *Rx3-Cre* that is constitutively active in the early optic vesicle starting at E8.75 and later in the optic cup in the presumptive retina, RPE and lens vesicle (77, 79, 80). Optic fissure (OF) closure starts at E10.5 and is completed by E12.5, evident by continuous pigmentation in the ventral optic cup at E13.5 (Fig. 1A, C). *Rx3-Cre*-mediated, conditional ablation of *Nf2* (hereafter *Nf2^CKO^*) results in failure of OF fusion in both eyes of each embryo (E13.5-E17.5; Fig. 1B, D; n = 22). In *Nf2^CKO^* optic cups, the OF margins are tightly apposed, suggesting a defect during the final process of closing (Fig. 1D). Persistent expression of laminin in the central ventral optic cup confirms failed degradation of the basement membrane between the OF margins (Fig. 1F; see Suppl. Fig. 1A for sagittal section scheme). We did not observe any coloboma in conditional heterozygous embryos (not shown). Laminin persistence in *Nf2^CKO^* shows 100% penetrance in the distal optic cup, close to the ciliary margin, while some can exhibit laminin degradation proximally at E13.5 (33%, 3/9 eyes, n = 7). Thus, coloboma in *Nf2^CKO^* eyes is less likely to be a delay in timing since failed closure in the distal and frequently in the proximal region of the optic cup is persistent at later stages. Compared to controls, the RPE layer in *Nf2^CKO^* eyes appears slightly darker (Fig. 1D, H; Suppl. Fig. 1C) and increased in height (Fig. 1G, H; insets). In addition, *Nf2^CKO^* embryos display additional, fully penetrant tissue fusion defects in closure of eye lids, lip and palate, as previously observed (Fig. 1B; not shown) (58).

**Figure 1.**
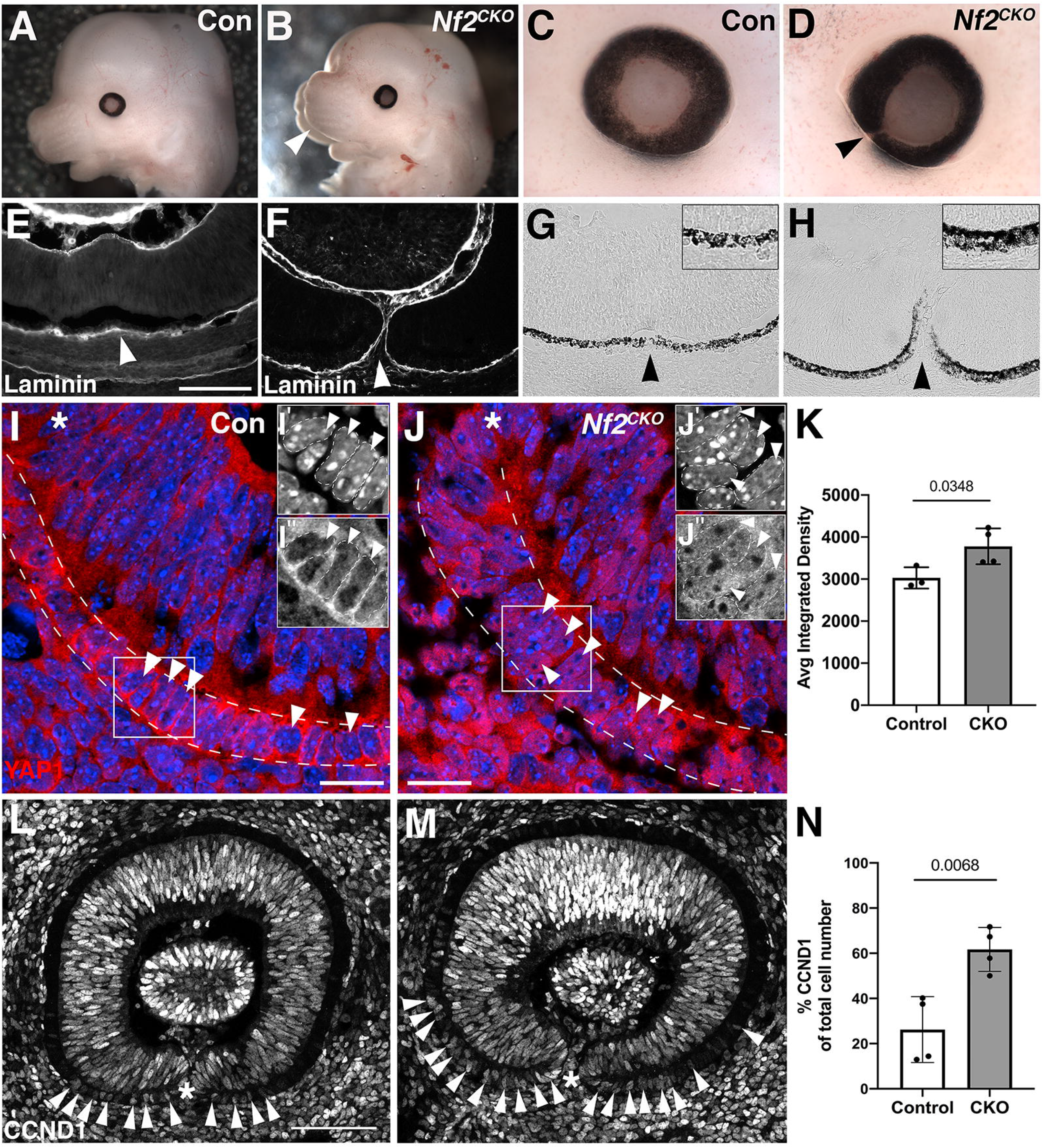
Nf2 disruption in ocular tissues causes coloboma in the embryonic mouse eye. A: Control embryonic head at E13.5. B: *Nf2^CKO^* embryos exhibit cleft lip and palate (arrowhead). C: Higher magnification of E13.5 control eye. D: In *Nf2^CKO^* embryos at E13.5, the ventral eye shows an OF closure defect with a narrow gap (arrowhead) and the pigment appears slightly darker. E: Immunolabeling for the basement marker laminin in the E13.5 ventral optic cup of controls confirms absence of laminin and successful fusion of the OF region (arrowhead). Sagittal view with nasal orientation to the left. F: Laminin is persistent in the ventral optic cup in E13.5 *Nf2^CKO^* embryos indicating failed OF fusion (arrowhead). G: Brightfield image of the ventral optic cup in control showing pigmentation of the RPE in the fused OF (arrowhead). H: In the ventral optic cup of E13.5 *Nf2^CKO^* embryos, pigmentation extends into the unfused OF (arrowhead) and the RPE layer appears slightly increased in height. Insets in G, H show magnified regions outlined in boxed area, revealing increased thickness in mutant RPE (H). I: During OF closure at E11.5, YAP1 protein expression can be localized both to the cytoplasm and nucleus in the RPE (arrowheads point to RPE nuclei). J: In *Nf2^CKO^* eyes, nuclear YAP1 expression can be increased in retina and RPE (arrowheads display nuclear Yap localization in RPE). RPE in I and J is indicated by dotted lines and sections were counterstained with Dapi (blue). Insets in I and J show enlarged greyscale images of individual cells in the RPE (boxed area). I’ and J’ display the Dapi channel, I’’ and J’’ display the YAP1 channel. Individual nuclei in the insets are outlined by dotted lines. K: YAP1 average integrated density per RPE nuclei shows a significant increase in *Nf2^CKO^* compared to control nuclei. L, M: Cyclin D1 (CCND1)-labeled cells are significantly increased in *Nf2^CKO^* ventral RPE (M; arrowheads; N). Asterisks in I, J, L, M indicate OF. Data are shown as means ± s.d., and p values are indicated above the horizontal lines in each graph (significant < 0.05). Statistical analysis was performed using unpaired, two-tailed Student T-test (Welch’s correction included for K). Scale bars: 100 μm in E, L; 20 μm in I, J.

A 2.4kB human *NF2* promoter element was recently identified to generate a *NF2-lacZ* reporter mouse, showing robust activation specifically in the RPE between E10.5 and E12.5 (81). NF2 protein is transiently expressed in the retina and continuously present in RPE and ciliary margin (59, 75). *Nf2* reporter activation and NF2 protein expression in the eye matches expression of the Hippo pathway target YAP. YAP1 protein normally shows cytoplasmic and nuclear localization in developing ocular tissues and in adjacent periocular mesenchyme (54, 59, 68, 72, 82, 83). Consistent with previous studies, we observed YAP1 expression in the cytoplasm and nucleus in retina and RPE during OF closure (Fig. 1I, insets I’-I” Suppl. Fig. 1F, M). In *Nf2^CKO^* optic cups, nuclear localization of YAP can be increased in cells of the RPE, as well as in retina and lens (Fig. 1J, insets J’-J”; Suppl. Fig. 1I, N; E11.0-E11.5; n=9). We quantified nuclear YAP signal intensity, confirming a significant increase in RPE cells adjacent to the OF as well as in the retina of *Nf2^CKO^* compared to controls (Fig. 1K; Suppl. Fig. 1K). Additionally, we calculated area of nuclei and observed no significant difference in the RPE and retina of control and *Nf2^CKO^* in E11.5 optic cups (Suppl. Fig. 1J, L). The ventral RPE layer in *Nf2^CKO^* can appear increased in thickness, possibly due to an increase in cell number and cellular crowding (Suppl. Fig. 1G, H, I; see also below Fig. 4). In addition, we investigated expression the cell cycle regulatory protein cyclin D1, a direct transcriptional target of YAP/TEAD in other tissues (84). We observed that in mutant RPE, significantly more cells express cyclin D1 in the ventral optic cup (Fig. 1M, N). Thus, our data strongly suggest that Nf2 function is required for retention of (phosphorylated) YAP in the cytoplasm during OF closure.

### In the ventral optic cup of *Nf2^CKO^*, RPE markers persist in the OF

Loss of *Nf2* results in increased nuclear YAP localization (Fig. 1), and YAP is required and sufficient to control RPE cell fate in zebrafish and mouse (67, 68). Therefore, we analyzed retina and RPE markers in the OF region between E11.5 and E13.5, during and after OF alignment and fusion, respectively (Fig. 2). The expression patterns of the pan-ocular paired PAX6 transcription factor (n=3) and retina-specific VSX2 homeodomain transcription factor (n=4) are unchanged in the ventral optic cup of *Nf2^CKO^* embryos at E11.5 (Fig. 2B, D). Dorsal localization of the T-box transcription factor TBX5 is unaltered in the *Nf2^CKO^* optic cup (Fig. 2F; n=4, E11.5-E13.5) (85). Similarly, ventral expression of the paired homeodomain transcription factor PAX2, essential for OF closure, is robust in the OF margins in *Nf2^CKO^* (Fig. 2H). Therefore, general tissue patterning of retina and dorsoventral regionalization is not affected by loss of *Nf2* function.

**Figure 2.**
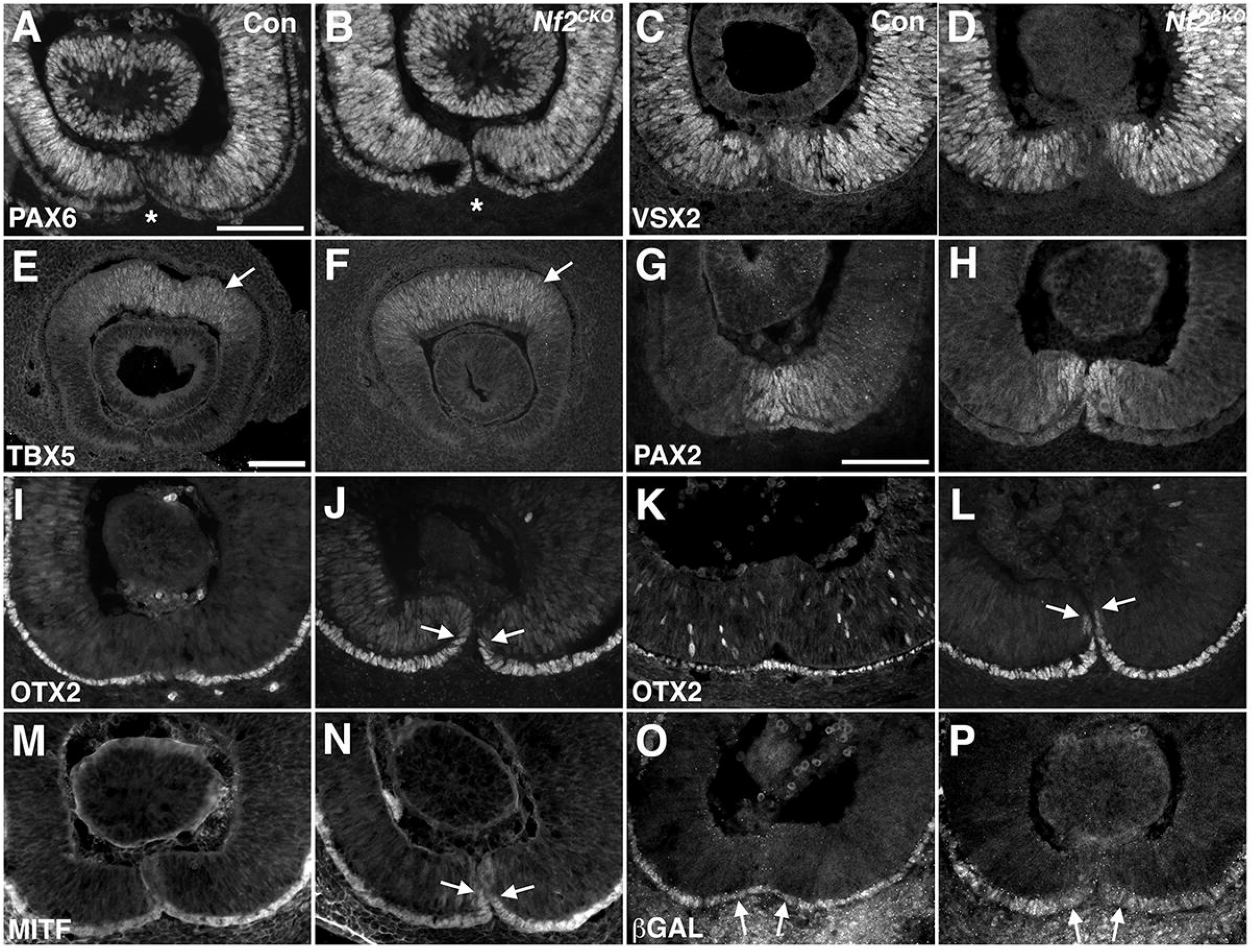
Analysis of tissue patterning in the *Nf2^CKO^* optic cup reveals persistence of RPE markers in the open optic fissure. (A-J, M-P: E11.5; K, L: E13.5) A, B: Expression of the pan-ocular protein PAX6 is unaltered in *Nf2^CKO^* eyes (B). Asterisks label overlying OF. C, D: The retina-specific homeodomain protein VSX2 is normally expressed in *Nf2^CKO^* eyes (D). E, F: Analysis of TBX5 expression shows normal patterning of the dorsal optic cup in *Nf2^CKO^* embryos (F; arrow). G, H: In the ventral *Nf2^CKO^* optic cup, the key transcription factor PAX2 is normally localized (H). I, J: Expression of the RPE determination transcription factor OTX2 can be maintained in the open fissure of *Nf2^CKO^* eyes (J, arrows). K, L: At E13.5, coloboma is persistent in *Nf2^CKO^* embryos and the margins of the unclosed OF are lined with OTX2-positive cells (L, arrows). M, N: The bHLH transcription factor MITF is present in the RPE layer of the optic cup (M). In *Nf2^CKO^* embryos, MITF localization extends into the OF (N; arrows). O, P: Axin2 reporter expression, a readout for Wnt/β-catenin pathway activation, is not altered in the ventral RPE in *Nf2^CKO^* optic cups (arrows). Sagittal view with nasal orientation to the left. Scale bars: 100 μm in A, E, G.

The bicoid transcription factor Otx2 is required for RPE cell fate and RPE differentiation at the advanced optic vesicle stage (86, 87). We observed that expression of OTX2 is ectopically maintained in the OF in several E11.5 *Nf2^CKO^* embryos (Fig. 2J; arrowheads; 57%; n=7). At E13.5 all analyzed eyes showed persistent OTX2 expression in the OF (Fig. 2L; arrowheads; 100%; n=4), consistent with extended pigmentation into the OF (Fig. 1D, H; Suppl Fig. 1C). The bHLH transcription factor MITF, a critical key regulatory factor for RPE cell fate and OF closure (18, 88, 89) is normally present in the approaching OF margins at E11.0 (Suppl Fig. 2A; arrowheads) and becomes progressively excluded ventrally during closing at E11.5 (Fig. 2M). At E11.0, MITF protein shows normal localization in the mutant OF (Suppl Fig. 2B; n=4). However, at E11.5, MITF-labeled cells extend into the lower (ventral) OF in *Nf2^CKO^* optic cups (Fig. 2N; arrows; 100%; n=4), suggesting persistent presence of RPE precursors in the OF. Abnormally upregulated RPE fate in the OF due to FGF pathway deficiency leads to failed OF fusion and YAP can cooperate with Pax transcription factors to promote MITF expression in neural crest progenitors (90–92). Therefore, to test whether excessive *Mitf* dosage in the OF margins is preventing OF closure, we removed one functional *Mitf* allele by crossing the *Mitf^Mi^* allele into *Nf2^CKO^* (89). Normally, heterozygous *Mitf^Mi/+^* optic cups are less pigmented (Suppl. Fig. 2D). In *Nf2^CKO^;Mitf^Mi/+^* optic cups, pigmentation is strongly reduced at E13.5, however, coloboma is still apparent (Suppl Fig. 2F, H; n=2). Thus, loss of *Nf2* function can result in maintenance of RPE cell fate in the OF margins, however, it may not be the primary cause for defective OF closure in *Nf2^CKO^* eyes.

### Wnt/β-catenin signaling and apicobasal polarity are unaltered upon loss of Nf2

Others and we have shown that Wnt/β-catenin signaling is required and sufficient for MITF and OTX2 expression in the mouse optic cup, and RPE-specific β-catenin gene disruption causes coloboma (93–96). However, gain of function of the pathway by disruption of the Wnt/β-catenin inhibitor *Axin2* can also result in coloboma, with persistent expression of MITF and OTX2 (19) similarly to *Nf2^CKO^*. Importantly, several studies have shown that NF2 can associate with LRP6 and block its phosphorylation resulting in inhibition of Wnt/β-catenin pathway activation (97). To determine whether Wnt/β-catenin pathway activity is hyperactivated upon loss of Nf2 function, we utilized a knock-in *Axin2^LacZ^* reporter (98). We did not detect obvious changes in *Axin2^LacZ^* (β-GAL) expression in *Nf2^CKO^* heterozygous for *Axin2^LacZ^* (Fig. 2P; n = 5). Additionally, expression of the downstream effector and target LEF1 as an additional readout for pathway activity was not altered (Suppl Fig. 2I, J; n = 4). These data indicate that Wnt/β-catenin pathway activity is not restrained by *Nf2* function.

NF2 participates in assembly of plasma membrane protein complexes and couples them with the cortical actin cytoskeleton (58, 60, 61, 63). In medaka, reduction of YAP results in decreased tension of the actomyosin network in embryonic tissues (99). Therefore, we reasoned that increased nuclear YAP may promote actomyosin activity in *Nf2^CKO^* ventral optic cups. Phosphorylated myosin light chain II (pMLC2) is normally localized to the apical RPE surface in the optic cup (Fig. 3A) (95, 100). In the mutant ventral RPE, adjacent to the OF, we observed no obvious changes in apical localization of pMLC2 protein (Fig. 3B; n=3). Similarly, localization of ZO-1, indicative for intercellular apical junction formation between retina and RPE, is not altered in *Nf2^CKO^* (Fig. 3D; n=5). NF2 also regulates assembly of adherens junctions (AJ) in the mouse epidermis, linking it to polarity protein complexes (63). However, we observed no change in the distribution of the AJ core protein α-catenin in the OF and apical surfaces (Fig. 3E, F; n = 3). In addition, F-Actin localization in the ventral optic cup of *Nf2^CKO^* appears normal (Fig. 6B, D; E11.0-E11.5; n=7). Thus, loss of *Nf2* in the optic vesicle does not interfere with apicobasal polarity, formation of adherens/tight junctions or actomyosin organization in the ventral optic cup during closure of the OF.

**Figure 3.**
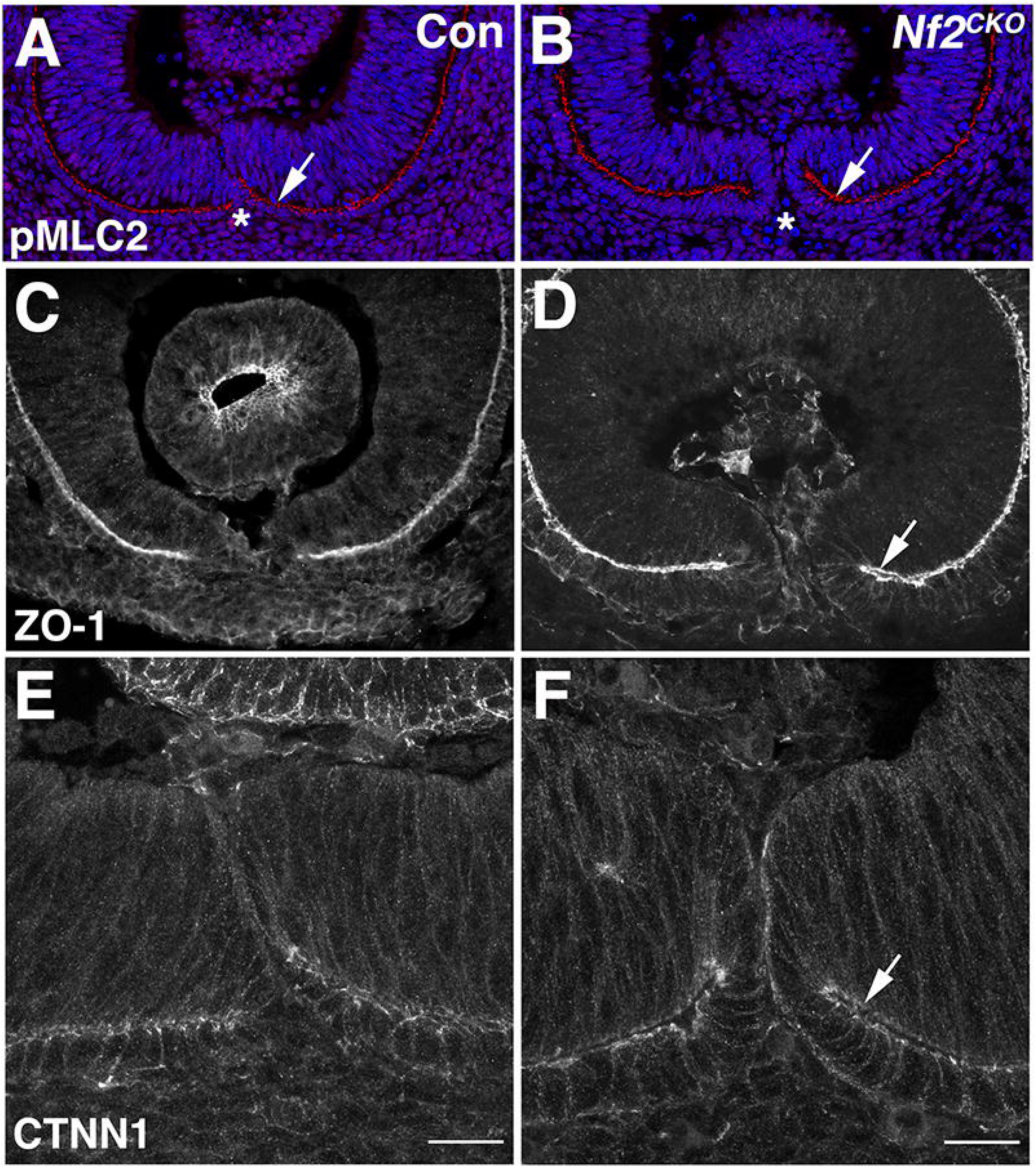
Apicobasal polarity and intercellular junction assembly are maintained during OF closure in the ventral optic cup in *Nf2^CKO^* embryos. (A, B, E, F: E11.5; C, D: E11.0) A, B: Localization of pMLC2 on the apical surface of the RPE layer appears normal in *Nf2^CKO^* embryos (B; arrow). Asterisk marks the overlying OF. C, D: Apical distribution of the tight junction component ZO-1 is not affected in the *Nf2^CKO^* ventral optic cup (D; arrow). The cadherin associated protein α-catenin shows normal localization in the OF and apical space between retina and RPE in *Nf2^CKO^* eyes (F, arrows). Sagittal view with nasal orientation to the left. Scale bars: 20 μm in E, F.

### During OF closure, ventral RPE proliferation is abnormally increased in *Nf2^CKO^* eyes

Depending on the species, OF closure is tightly regulated by a balance of programmed cell death (apoptosis) and proliferation. In zebrafish, apoptosis is not abundant in the ventral optic cup, and cell death or proliferation are not required for OF closure (27, 34, 46, 101, 102). In chick and mammals, retinal apoptosis is prevalent and retinal proliferation is decreased in the vicinity of the OF margins during closure (9, 10, 27, 103–105). Accordingly, abnormal changes in cell number by altered cell death or cell divisions in the optic cup can result in OF closure defects and colobomata (44, 106). Since *Nf2* disruption causes altered apoptosis during neural tube closure (58), we first analyzed programmed cell death using TUNEL labeling. In the ventral optic cup of E11.5 *Nf2^CKO^* embryos, apoptosis was not significantly altered in the ventral quadrant harboring the OF (V2; Suppl. Fig. 3A-D; see Suppl. Fig. 3C for subdivision scheme).

Nuclear localization of YAP can stimulate cell division, and a significant increase in RPE marker-labeled cells and BrdU incorporation in the RPE layer is observed in *Nf2^CKO^* using *Nestin-Cre* (58, 59). However, spatial and temporal information critical for the process of OF closure is lacking; these studies were performed after completion of OF closure, and not regionally correlated with the OF. Thus, we performed a detailed proliferation analysis in *Nf2^CKO^* optic cups during OF closure. Phospho-histone H3-labeling to detect proliferating cells in G2/M of the cell cycle revealed an upward trend in the ventral *Nf2^CKO^* optic cup that was not significant (Fig. 4A-C; E11.5; n=3). However, we observed a small but significant increase of phosphohistone H3-labeled cells in the ventral quadrant that includes the OF region (Fig. 4D). Since the number of G2/M cells, particularly in the RPE, was generally very low in the optic cup, we additionally analyzed EdU incorporation (Fig. 4E-J). We determined changes in proliferation specifically during the initial alignment of the OF margins, a critical stage for the subsequent attachment and fusion processes during OF closure (E11.0; n=4). The percentage of EdU-labeled cells is significantly increased by 46% in the *Nf2^CKO^* RPE (Fig. 4F, G). In contrast, no significant change is detectable in the retina (Fig. 4G, H). The RPE-specific increase in EdU-labeled cell number is specific for the ventral portion in *Nf2^CKO^* optic cups, by 79% (Fig. 4F, I; V1-V3), while the dorsal RPE shows no significant change at this stage (Fig. 4F, I). Significantly higher numbers of EdU-labeled RPE cells are observed in all subdivisions of the ventral optic cup, including the optic fissure region (V2; Fig. 4J). Consistent with stimulated proliferation, the total number of cells in the ventral optic cup is significantly expanded in *Nf2^CKO^* RPE (Suppl. Fig. 3E). In summary, our data indicates that Nf2 disruption in the optic vesicle results in abnormally increased RPE proliferation in the ventral optic cup early in the process of OF closure.

**Figure 4.**
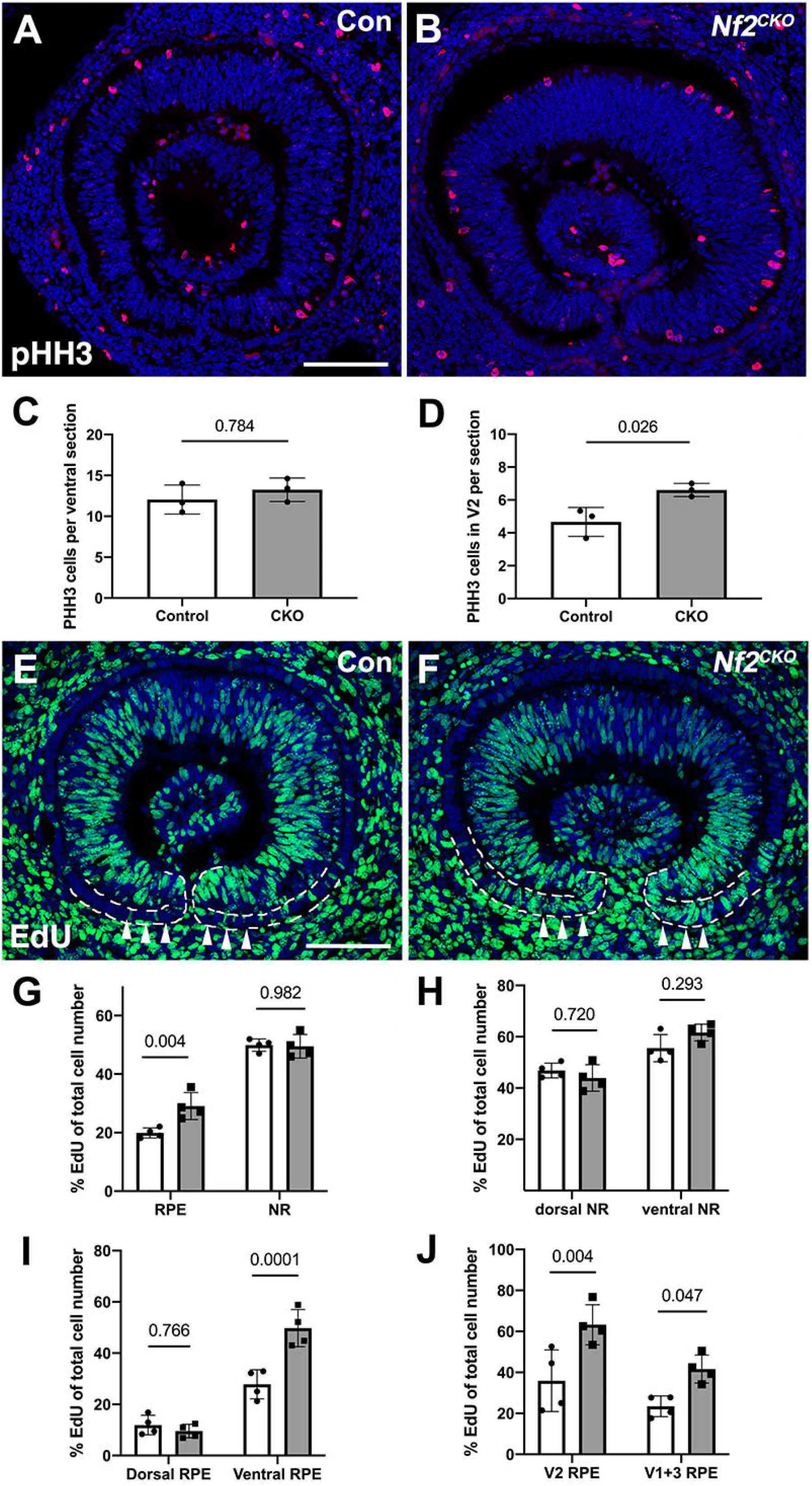
Proliferation is upregulated in the RPE layer in the *Nf2^CKO^* optic cup. (A-D: E11.5; E-J: E11.0) A-B: Phospho-histone H3 (pHH3) labeling (red) of control (A) and *Nf2^CKO^* (B) detects ocular cells in G2/M phases of the cell cycle (red). Dapi is used as a nuclear label (blue). Quantification shows that the number of phospho-histone H3-labeled cells is not changed in the ventral optic cup in *Nf2^CKO^* eyes (C); however, there is a small significant increase in the ventral-most area that harbors the OF (V2; D). E-J: EdU labeling and quantification. EdU-labeled cells in control (E; green), with nuclear Dapi co-labeling (blue). In the RPE of the *Nf2^CKO^* ventral optic cup, the number of EdU-labeled cells is increased (F; arrowheads), in comparison to controls (E, arrowheads). Quantitative analysis of EdU incorporation reveals that the number of cells in S-phase is significantly upregulated specifically in the RPE but not in the retina (G, H). (RPE: control: 19.91% ± 1.74 s.d.; *Nf2^CKO^*: 29.07% ± 4.60 s.d.) The increase in EdU-labeled cell number is significant for the ventral RPE (I; control: 27.82% ± 5.66 s.d.; *Nf2^CKO^*: 49.86% ± 7.10 s.d.), while the dorsal RPE shows no change. Significant stimulation of proliferation in the ventral RPE applies to all regions in the ventral optic cup, including the V2 region harboring the OF (J). Data are shown as means ± s.d., and p values are indicated above the horizontal lines in each graph (significant < 0.05). Statistical analysis was performed using unpaired, two-tailed Student T-test or 2-Way ANOVA with Sidak’s multiple comparison. Images show sagittal view with nasal orientation to the left. Scale bar: 100 μm in A, E.

### Ectopic RPE cell number in the dorsal optic cup of *Nf2^CKO^* embryos

As described above, we detected no significant increase in the percentage of EdU incorporation in the dorsal *Nf2^CKO^* RPE (Fig. 4I). However, we observed supernumerary cells and frequent cellular crowding in the MITF-labeled RPE layer of the dorsal optic cup (Fig. 5B). Quantification of total cell number revealed a robust significant increase by 47% specifically in the dorsal *Nf2^CKO^* RPE layer (Fig. 5C; n = 4). In contrast, total cell number in the dorsal retina did not change (Fig. 5D). Furthermore, we observed regions with irregular F-actin organization and mis-localization of pMLC2 in the dorsal RPE layer (Fig. 5F, H; arrowheads). Abnormally increased number of RPE cells with disorganization of the RPE layer was maintained later at E13.5, using OTX2 as a marker for differentiating RPE cells (Fig. 5J; arrowheads). Collectively, our data indicates that ocular *Nf2* disruption leads to a significant increase of cell number in the dorsal and ventral RPE layer of the optic cup. This occurs with concomitant acquisition of RPE cell fate and mis-localization of F-Actin and pMLC2. However, size of the *Nf2^CKO^* eye is unchanged, since measurement of the ocular diameter in embryos did not reveal a significant difference (Suppl. Fig. 3F).

**Figure 5.**
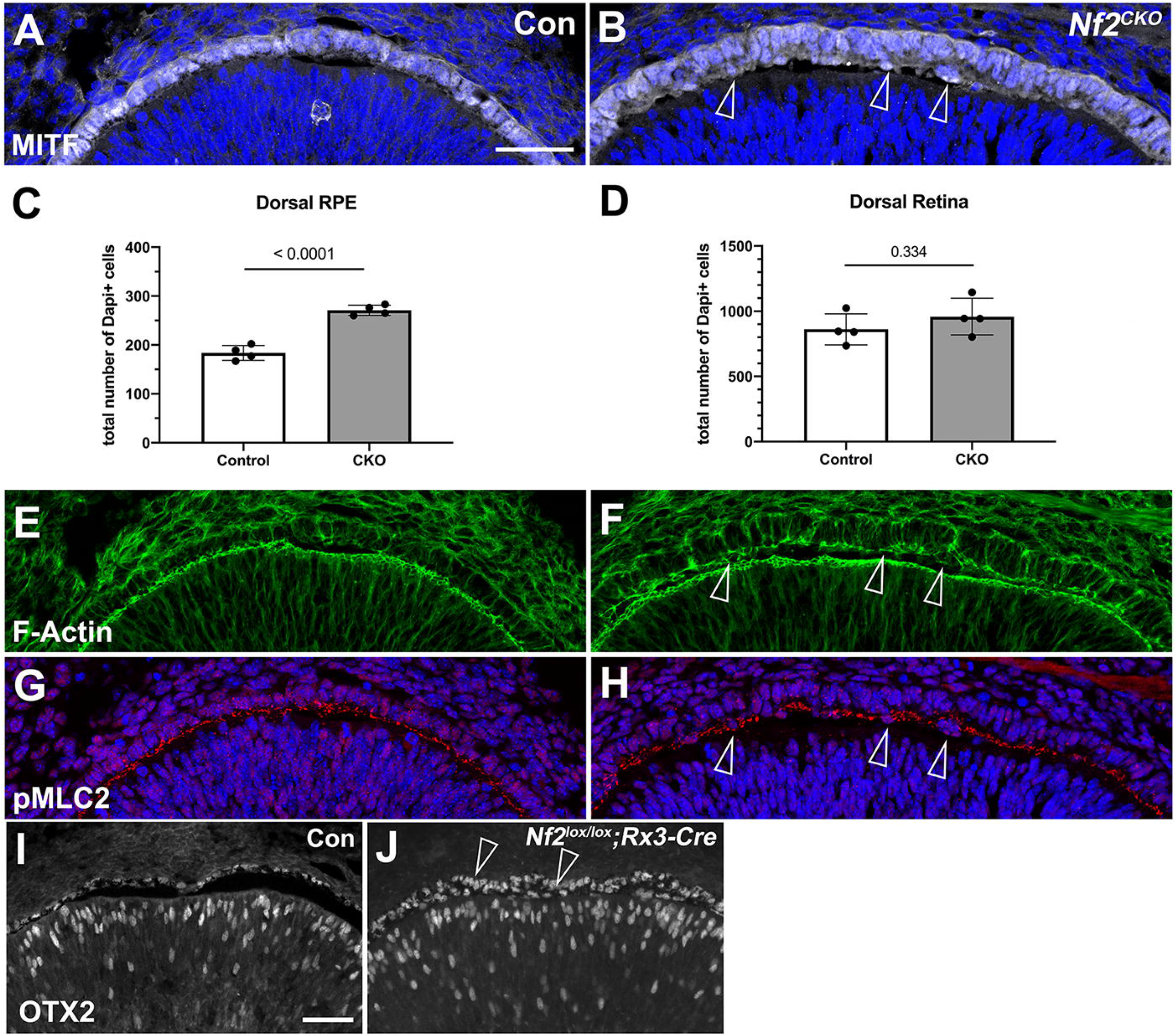
Cell number in the RPE layer is increased in the *Nf2^CKO^* dorsal optic cup at E11.0. A, B: MITF labeling reveals increased RPE layer thickness in *Nf2^CKO^* (B; arrowheads point to ectopic RPE cells located apically). C, D: Quantification indicates a significant increase in total cell number in the dorsal RPE in *Nf2^CKO^* (C; see Suppl. Fig. 3C for subdivisions; control: 183.75 ± 15.13 s.d.; *Nf2^CKO^*: 271.00 ± 10.42 s.d.), while cell number is unchanged in the dorsal retina (D). Data are means ± s.d., Student’s T-test was applied for statistical analysis, and p values are indicated on the horizontal lines in each graph (significance < 0.05). F: Phalloidin-labeling reveals irregularities in localization in the dorsal RPE layer in *Nf2^CKO^* (arrowheads). H: In addition, mislocalization of pMLC2 is detectable in the dorsal RPE layer in *Nf2^CKO^* optic cups (arrowheads). I, J: OTX2 labeling shows that supernumerary cells persist in the dorsal optic cup at E13.5 (J; arrowheads). Scale bars: 50 μm in A, I.

### Alignment and cellular height are altered in the OF margins of the *Nf2^CKO^* optic cup

Early in the process of OF closure in controls, the fissure margins become tightly aligned, shaped with a convex surface toward the basal space (Fig. 6A). In the *Nf2^CKO^* OF, the shape of the basal surface of the OF margins is convex, similar to controls, however, alignment of the margins is less advanced (Fig. 6B). At E11.5, margins in control embryos are well aligned along most of the OF with a flattened basal surface (Fig. 6C). In E11.5 *Nf2^CKO^* optic cups, we noticed that the shape of the approaching OF margins is consistently altered; the margins maintain a convex shape and the actual extent of contact area between both OF margins is often reduced (Fig. 6D). We quantified this by measuring the extent by which the margins become tightly apposed and determined that the OF in *Nf2^CKO^* eyes shows a significant reduction (Fig. 6E).

**Figure 6.**
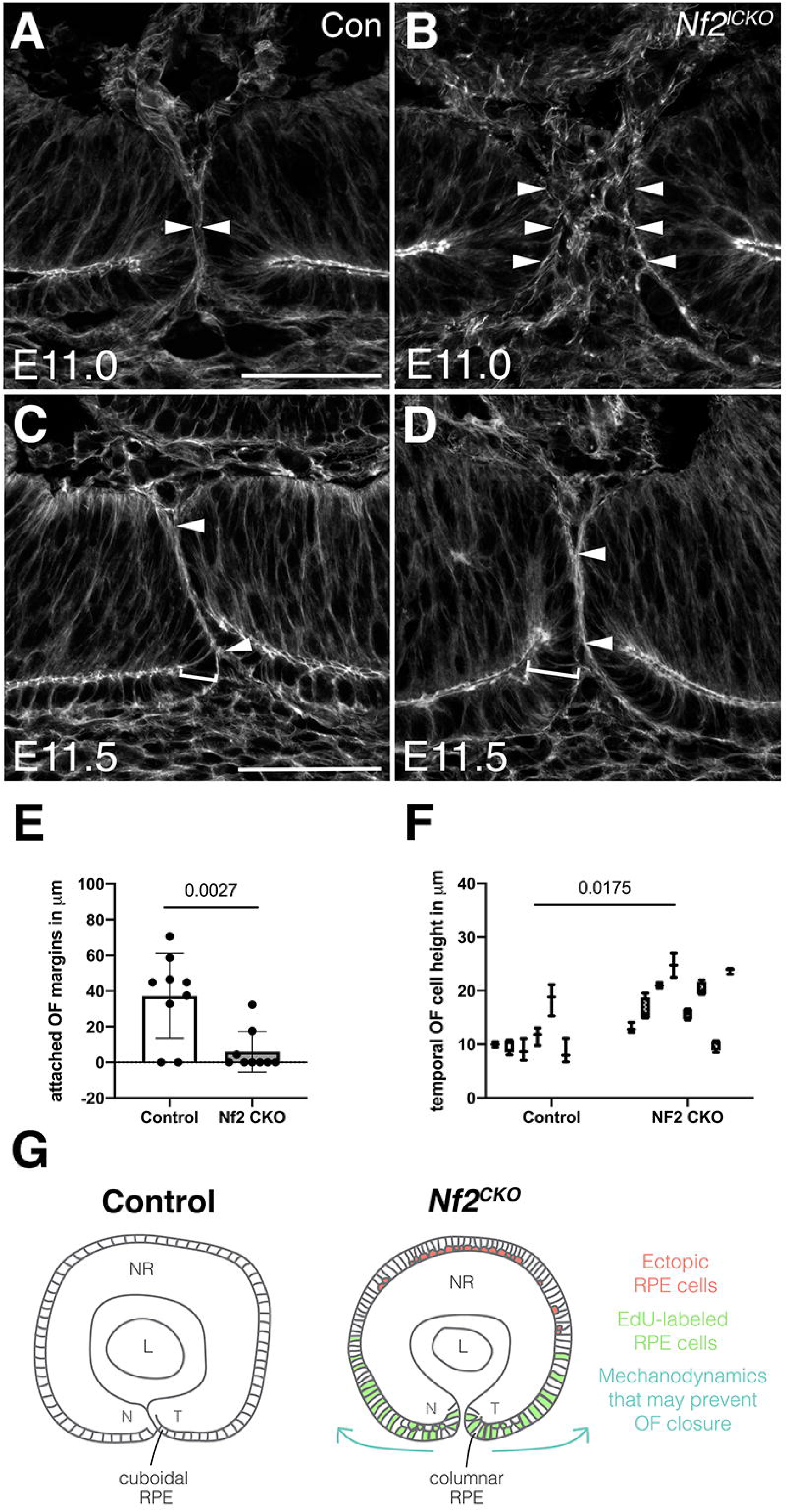
*Rx3-Cre*-mediated disruption of Nf2 results in defective alignment of OF margins during the closure process. A-D: F-actin localization in the OF at E11.0 (A, B) and at E11.5 (C, D) in control (A, C) and *Nf2^CKO^* optic cups (B, D). A: In E11.0 control embryos, the OF margins become tightly aligned, shaped with a convex surface toward the basal space (arrowheads). B: In the *Nf2^CKO^* OF, shape of the basal surface of the OF margins is convex, similar to controls, however, alignment of the margins is less advanced (B, arrowheads). At E11.5, the OF margins in controls are well aligned along most of the OF with flattened basal surfaces (C, arrowheads). The RPE layer consists of cuboidal cells (bracket). D: In *Nf2^CKO^* eyes, the OF margins align only partially (arrowheads), exhibit convex shape, and OF cells continuous with the RPE layer show increased height in the apicobasal axis, thereby appearing columnar (bracket). E: Quantification of aligned OF margins. Extent of attached margins was measured in the OF, as outlined in representative examples shown in C, D for aligned OF between arrowheads. In *Nf2^CKO^*, few margins show close alignment, which is often much reduced, compared to controls (lack of alignment defined = 0). Data are means ± s.d. and Student’s T-test was applied for statistical analysis. F: Analysis of cellular height along the apicobasal axis of cells lining the ventral OF margins (see brackets in C, D; Suppl. Fig. 3G) reveals significantly increased cellular height in the temporal OF margin. Statistical analysis was performed using nested T-test, data are means (with minimal and maximal points). P values are indicated on the horizontal lines in each graph (E, F). G: Proposed model summarizing our observations, for explanation, see text (left: control optic cup; right: *Nf2^CKO^*). The arrows point to potential effects of RPE hyperproliferation on the mechanodynamics of optic cup growth and apposition of OF margins. Sagittal view with nasal orientation to the left. (NR: neural retina, L: lens, N: nasal, T: temporal) Scale bars: 50 μm in A, C.

We observed a tighter cellular packing and crowding with occasional formation of a double layer in *Nf2^CKO^* RPE (Fig. 4F). This is most likely caused by increased proliferation particularly in the ventral RPE layer that is continuous with the ventral OF margins (V2; Fig. 4J). In addition, we found that the ventral marginal OF layers appears increased in height along the apicobasal axis and columnar in shape, in comparison to the rather cuboidal organization in controls (Fig. 6C, D; brackets). We measured the apicobasal height of cells lining each temporal and nasal margin in the ventral OF (Suppl. Fig. 3G) and observed a significant increase in cellular height particularly in the temporal OF in *Nf2^CKO^* (Fig. 6F). In the nasal OF, cell height measurement showed an upward trend that is not significant (Suppl. Fig. 3H). These results suggest that, in the absence of *Nf2* function, convex shape of the OF margins is maintained, combined with a columnar organization, resulting in insufficient alignment and subsequent failure to fuse.

## Discussion

In summary, our results demonstrate that conditional, ocular disruption of the *Nf2* gene at the optic vesicle stage results in a fully penetrant, distally persistent late-stage defect in OF closure in the mouse embryo (see Fig. 6G for a proposed summary). *Nf2^CKO^* eyes show increased nuclear localization of YAP protein in ocular tissues, without apparent defects in formation of junctional complexes or apicobasal polarity in the ventral optic cup. Here we identify expansion specifically of the RPE layer as a unique underlying cause for coloboma. In *Nf2^CKO^* mutant eyes, more cells in the RPE layer in the ventral optic cup, including in the OF margins, proliferate and acquire RPE fate. In the OF margins, cells extend in height or double up as a second layer, and the RPE layer maintains columnar organization instead of transitioning into cells of cuboidal shape. In addition, in the RPE of the dorsal optic cup, *Nf2* loss of function leads to a robust increase in cell number, with local disorganization of the cytoskeleton components F-actin and pMLC2. We propose that this specific effect on proliferation in the RPE layer in the *Nf2^CKO^* optic cup may result in changes in mechanodynamics within the growing optic cup, causing the OF margins to align insufficiently.

### Nf2 is required for balancing RPE cell number during OF closure

During optic cup morphogenesis, the RPE is required for growth and OF closure; extreme, broad removal of RPE by diphtheria toxin in the early optic cup causes microphthalmia and coloboma (29, 107). Growth of the RPE during eye development is dynamic; proliferation is high in the entire optic vesicle (50-60%) and decreases in the RPE substantially soon after invagination (12-25%) (108–111). During normal OF closure, cell number in the mammalian optic cup is tightly balanced by cell proliferation and programmed cell death (13, 44, 106). Particularly close to the approaching OF margins, proliferation is lower compared to other regions of the optic cup, and deregulated proliferation leads to coloboma (44). This is assumed to prevent cellular crowding in the fissure to allow for cellular reorganization. However, it is unclear from these studies whether cell number specifically in the RPE layer is critical for OF closure (44, 91, 106). *Nf2^CKO^* eyes exhibit a substantial increase in proliferation in the ventral RPE when the OF margins are approaching (by 79%; Fig. 4I), as well as a robust increase of overall cell number in the dorsal RPE. Furthermore, RPE cell expansion is only observed when *Nf2* loss occurs in the optic vesicle and early optic cup, but not at later time points (59)(SR and SF; unpublished observations). This is consistent with the inability of the RPE to renew itself, once differentiation and further maturation commences (112). Few factors have been identified that can promote proliferation and expansion of RPE; these include transgenic hyperactivation of Notch signaling, resulting in microphthalmia and coloboma (29, 113), or loss of factors that normally limit proliferation in the developing RPE, such as the cyclin-dependent kinase inhibitor p27Kip1 or DAPL1 (114–116). Collectively, our analysis reveals a unique role for *Nf2* in regulating closure of the optic fissure by restricting RPE proliferation in the optic cup.

### Ectopic cells in the outer layer of the *Nf2^CKO^* optic cup acquire RPE fate

In the growing optic cup, the presumptive RPE slowly proliferates to maintain a single layer and differentiates into pigmented cells in a central to peripheral and dorsal to ventral gradient, which is tightly coordinated with growth of the neural retina (88, 110, 111). Abnormal increases in proliferation in the presumptive RPE in the early optic cup is usually induced by loss of RPE key regulatory gene function or by direct misexpression of factors stimulating proliferation, for example, by modulation of extracellular matrix molecules, extracellular signaling pathways (e.g. FGF, Wnt, Shh), overexpression of oncogenes and loss of key RPE determination genes (112, 117). Since the presumptive RPE is bipotential at the early optic cup stage, these manipulations result in transdifferentiation of RPE into retina, with concomitant acquisition of neural retina-specific cell fate and formation of an ectopic retina (for review, see (112)). Our findings are unusual; in *Nf2^CKO^* eyes, proliferation of the presumptive RPE is strongly upregulated without inducing transdifferentiation into retina, and more RPE cells are produced. This effect is associated with an increase in nuclear YAP localization, consistent with YAP/TAZ’s role in regulating RPE cell fate and differentiation in zebrafish and mouse (59, 66–68). In addition, YAP mutant eyes develop coloboma in zebrafish. Thus, YAP/TAZ likely exerts a dual role in promoting proliferation and differentiation in the presumptive RPE layer in *Nf2^CKO^* optic cups. In premigratory neural crest progenitors, YAP cooperates with PAX3 to transactivate *Mitf*, thus, YAP/TAZ may directly regulate RPE-specific gene expression (92).To further confirm that YAP/TAZ are required for RPE expansion in *Nf2^CKO^* embryos, we attempted to block YAP/TEAD interaction by administering Verteporfin; however, this was not successful, possibly due to solubility issues. A recent analysis in mouse revealed that NF2 coordinates later proliferation and differentiation of peripheral retina, ciliary body/iris and RPE by limiting nuclear YAP/TAZ activation (59). Thus, it is very likely that RPE expansion in *Nf2^CKO^* eyes is mediated via nuclear translocation of YAP/TAZ and activation of TEAD.

### Coloboma may be due to persistent buckling of the OF margins

In *Nf2^CKO^* optic cups apposition of both OF margins is significantly decreased; the margins show a higher degree of bending and do not flatten as observed in controls during OF alignment (Fig. 6). Although, some optic cups can exbibit fusion at varying degrees proximally close to the optic disc, the OF closure defect in *Nf2^CKO^* optic cups is always persistent in the distal optic cup at later stages, excluding a simple delay. While we cannot rule out other potential mechanisms (e.g. constraints provided by the extracellular matrix), our results suggest that over-proliferation in *Nf2^CKO^* RPE is likely to be the primary defect. Expanded proliferation may exert a critical mechanical force for promoting tissue buckling, thereby possibly resulting in prolonged convex bending of the OF margins. In a potentially similar scenario, experimentally induced, localized increases in proliferation due to constitutively activated β-catenin can exert mechanical forces to stimulate cortical folding in the embryonic mouse telencephalon (118). Thus, it is possible that similar effects occur on mechanodynamics in the *Nf2^CKO^* optic cup.

Our observations suggest that shape and organization of the RPE layer may be important for OF closure. RPE morphogenesis during formation of the optic cup is dynamic; RPE cells transition from a pseudostratified columnar epithelium into a single layer of cuboidal cells (for reviews, see (87, 88)). Interestingly, in zebrafish, RPE cells flatten further and expand around the retina in the optic cup, independent of cell proliferation and possibly by recruitment of additional cells into the layer (46, 119) (for review, see(120)). Electron microscopic studies in mouse and hamster suggest that marginal OF cells in the immediate vicinity of the first fusion contact re-orient their polarity during degradation of the basement membrane (9, 10). This is supported by live imaging studies in zebrafish, revealing extensive movements and incorporation of OF marginal cells into retina as well as RPE during closure (34, 49). In *Nf2^CKO^* RPE, cells in the temporal OF margin do not properly transition to a cuboidal monolayer at the time of closure (Fig. 6F). It is possible that the expansion of *Nf2^CKO^* RPE results in cellular packing/crowding, preventing cells from reorienting properly in the OF. This re-arrangement may be necessary to prepare for fusion at the right time. However, we cannot fully exclude that the cells lining the OF harbor a primary intrinsic defect that ultimately prevents proper dissolution of the basement membrane.

### Role of Hippo signaling in OF closure in humans

Hippo pathway components have been recently identified as genes with clinical relevance for coloboma (23, 53, 54, 64). Heterozygous loss of function mutations in YAP1 were identified in 2 unrelated families exhibiting coloboma (3). According to previous studies using animal models, YAP1 may be important for regulating RPE cell fate and RPE proliferation during OF closure in humans (67, 68). However, YAP1 is also present in the periocular mesenchyme (Fig. 1; (54)), which could be of importance for studies addressing the syndromic occurrence of coloboma. Furthermore, a mutation in the *Tead1* gene causes SCRA, preventing it from interacting with YAP1 (64, 121). SCRA manifests as bilateral stripes of atrophic retina and choroid; however, the contribution of RPE defects in disease pathology is not well understood. The protocadherin FAT1 represents a potential upstream regulator of the Hippo pathway, and it is expressed in the RPE (23). A new clinical syndrome with frequent ocular manifestations, in particular coloboma, is reportedly caused by frameshift mutations in the *FAT1* gene that may restrict nuclear localization of YAP. While FAT1 may be critical for cell-cell contact during OF closure, a direct effect on YAP localization was not observed (23). Furthermore, *NF2* mutations cause a variety of ocular defects particularly in the RPE; for example, increased pigmentation and combined retinal and RPE hamartomas (53, 122). Importantly, coloboma was observed in NF2 patients with germline nonsense mutations (53). Thus, it is possible that hyperproliferation of the RPE represents an underlying mechanistic cause for some of the observed ocular abnormalities, including coloboma, in individuals with inherited *NF2* mutations.

## Material and Methods

### Mice

Animal procedures were approved by the Vanderbilt University Medical Center Institutional and Animal Care and Use Committee. *Nf2^lox/lox^* mice (77) were crossed with *Rx3-Cre* (80), *Axin2^lacZ^* (Jax strain #009120); (98)), and *Mitf^Mi^* (Jax strain #001573 (123, 124)). Mouse lines were maintained in a mixed genetic C57BL/6 and CD-1 background. No ocular or extraocular defects were detected in *Nf2^lox/+^;Rx3-Cre* eyes, and littermates without *Cre* allele were used as controls. Some samples were heterozygous for the *Axin2^lacZ^* reporter allele, which did not reveal any difference to samples harboring the *Axin2^wt^* allele. Some control samples were heterozygous for the *Rd8* mutation, and no difference to *Rd8* wildtype controls was observed. The *Rd8* mutation results from a mutation in the gene *Crb1* (125, 126), however, only ocular abnormalities at postnatal ages such as retinal disorganization and photoreceptor degeneration have been observed (127). For timed pregnancies, counting started at day 0.5 when the vaginal plug was detected. Embryonic tissues were genotyped using the following primer combinations (19, 77, 79). The *Nf2* primers CTT CCC AGA CAA GCA GGG TTC (forward, P4) and GAA GGC AGC TTC CTT AAG TC (reverse, P5) generate amplicons for *wt* (305 bp) and *lox* (442 bp); *Rx3-Cre* primers: GTT GGG AGA ATG CTC CGT AA (forward) and GTA TCC CAC AAT TCC TTG CG (reverse) produce a 362 bp amplicon; and the *Axin2^lacZ^* primers: AAG CTG CGTCGG ATA CTT GAG A (Cs), AGT CCA TCT TCA TTC CGC CTA GC (Cwt), TGG TAA TGC TGC AGT GGC TTG (ClacZ) generate the *Axin^wt^* (493 bp) and *Axin^lacZ^* amplicons (400 bp). For genotyping of the *Mitf^Mi^* allele, tissue samples were processed and genotyped by Transnetyx (Cordova, TN) using Taqman with custom-designed primers: CCTTTCCCATGCTCTTTTCTTGAAG (forward), CTAGCTCCTTAATGCGGTCGTTTAT (reverse), and the following reporter probes to detect the 3-nucleotide mutation: ACGAAGAAGAAGATTTAAC (wildtype allele), AACGAAGAAGATTTAAC (*Mitf^Mi^* allele). To detect proliferating cells in S-phase of the cell cycle, pregnant dams received intraperitoneal EdU injections of 30 μg/g body weight 2 hours before sacrificing (Thermofisher/Invitrogen, Rockford, IL; #E10187).

### Immunohistochemistry

Embryos at the indicated ages were processed as previously described (79). Briefly, embryonic heads were fixed in 4% paraformaldehyde (PFA) for 1 hour, cryoprotected in sucrose, sectioned on a Leica CM1950 cryostat at 12 μm and stored at −80C until further use. For immunohistochemical analysis, sections were processed using antigen retrieval with triton X-100 (0.1%, 1%) and hot citrate buffer pH 6.0. Primary and secondary antibody information is provided in Table 1. Filamentous actin was detected using phalloidin (1:50; Thermo Fisher Scientific, Walham, MA; #A12379). Labeling with WGA-555 (1:200; Thermo Fisher Scientific, Walham, MA; #W32464) was performed prior to antigen retrieval. For detection of apoptotic cells, ApopTag Fluorescein In Situ Apoptosis Detection Kit (EMD Millipore, #S7110) was used according to the manufacturer’s instructions. EdU detection was performed using the Click-iT^®^ EdU Imaging Kit (Thermo Fisher Scientific, Walham, MA; #C10637). Sections were counterlabeled with DAPI (Thermo Fisher Scientific, Walham, MA; #D3571) and mounted in Prolong Gold Antifade. A minimum of 3 eyes from 3 different embryos from at least 2 individual litters were analyzed per genotype, time point and marker.

**Table 1:**
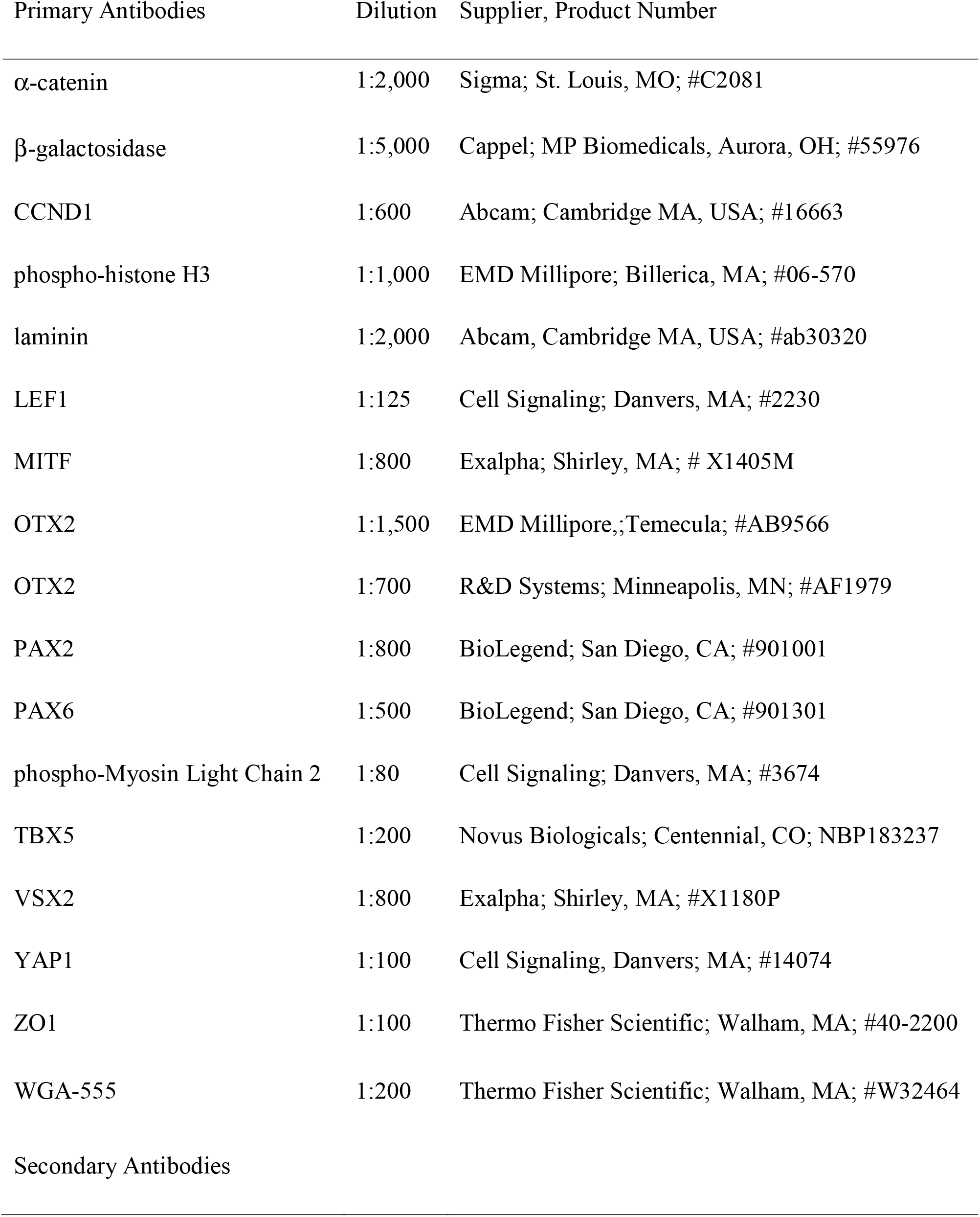
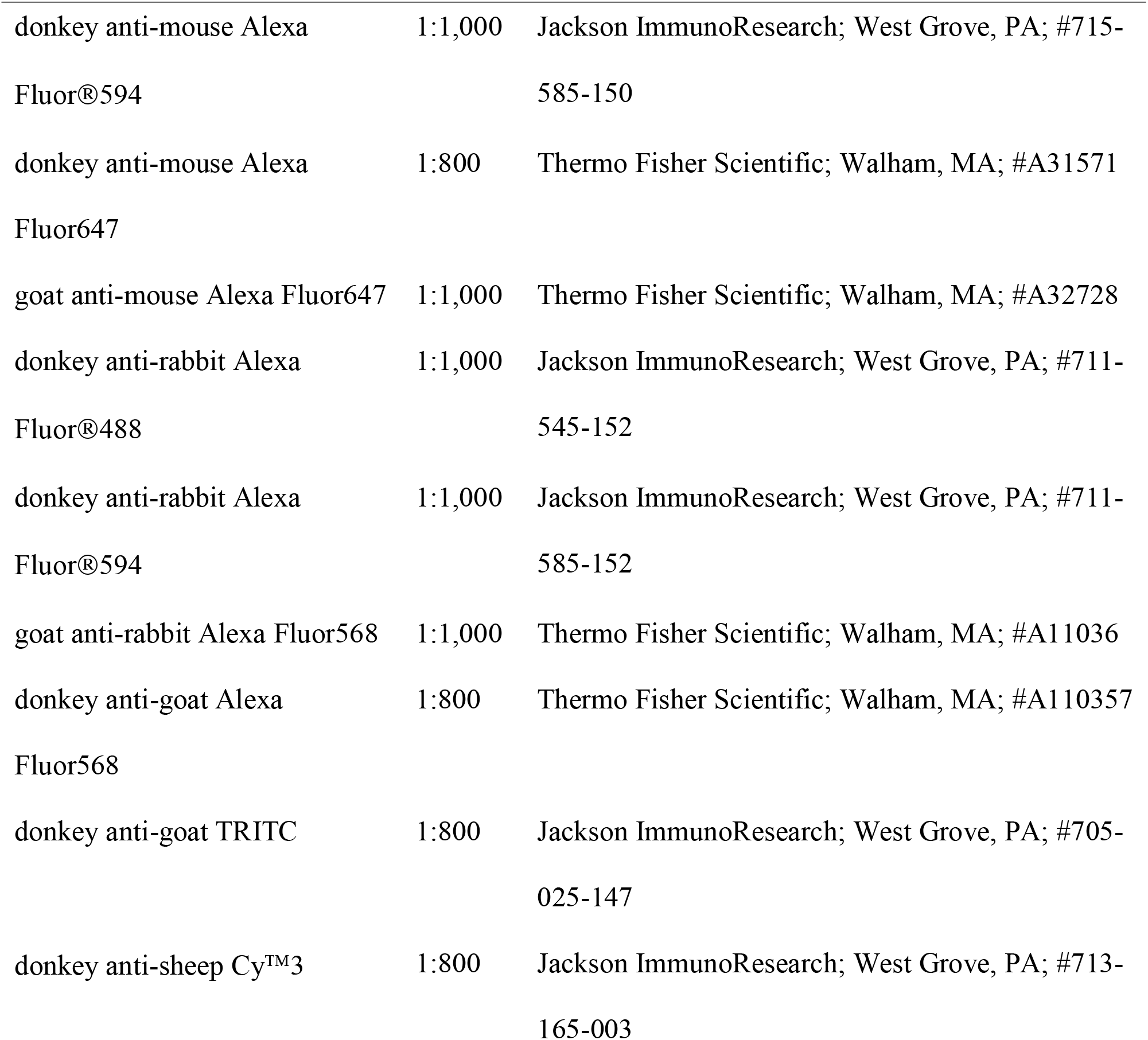
Antibodies used in this study

| Primary Antibodies | Dilution | Supplier, Product Number |
| --- | --- | --- |
| α-catenin | 1:2,000 | Sigma; St. Louis, MO; #C2081 |
| β-galactosidase | 1:5,000 | Cappel; MP Biomedicals, Aurora, OH; #55976 |
| CCND1 | 1:600 | Abcam; Cambridge MA, USA; #16663 |
| phospho-histone H3 | 1:1,000 | EMD Millipore; Billerica, MA; #06-570 |
| laminin | 1:2,000 | Abcam, Cambridge MA, USA; #ab30320 |
| LEF1 | 1:125 | Cell Signaling; Danvers, MA; #2230 |
| MITF | 1:800 | Exalpha; Shirley, MA; # X1405M |
| OTX2 | 1:1,500 | EMD Millipore,; Temecula; #AB9566 |
| OTX2 | 1:700 | R&D Systems; Minneapolis, MN; #AF1979 |
| PAX2 | 1:800 | BioLegend; San Diego, CA; #901001 |
| PAX6 | 1:500 | BioLegend; San Diego, CA; #901301 |
| phospho-Myosin Light Chain 2 | 1:80 | Cell Signaling; Danvers, MA; #3674 |
| TBX5 | 1:200 | Novus Biologicals; Centennial, CO; NBP183237 |
| VSX2 | 1:800 | Exalpha; Shirley, MA; #X1180P |
| YAP1 | 1:100 | Cell Signaling, Danvers; MA; #14074 |
| ZO1 | 1:100 | Thermo Fisher Scientific; Walham, MA; #40-2200 |
| WGA-555 | 1:200 | Thermo Fisher Scientific; Walham, MA; #W32464 |
| Secondary Antibodies |  |  |
| donkey anti-mouse Alexa Fluor®594 | 1:1,000 | Jackson ImmunoResearch; West Grove, PA; #715-585-150 |
| donkey anti-mouse Alexa Fluor647 | 1:800 | Thermo Fisher Scientific; Walham, MA; #A31571 |
| goat anti-mouse Alexa Fluor647 | 1:1,000 | Thermo Fisher Scientific; Walham, MA; #A32728 |
| donkey anti-rabbit Alexa Fluor®488 | 1:1,000 | Jackson ImmunoResearch; West Grove, PA; #711-545-152 |
| donkey anti-rabbit Alexa Fluor®594 | 1:1,000 | Jackson ImmunoResearch; West Grove, PA; #711-585-152 |
| goat anti-rabbit Alexa Fluor568 | 1:1,000 | Thermo Fisher Scientific; Walham, MA; #A11036 |
| donkey anti-goat Alexa Fluor568 | 1:800 | Thermo Fisher Scientific; Walham, MA; #A110357 |
| donkey anti-goat TRITC | 1:800 | Jackson ImmunoResearch; West Grove, PA; #705-025-147 |
| donkey anti-sheep Cy™3 | 1:800 | Jackson ImmunoResearch; West Grove, PA; #713-165-003 |

Embryo and whole eye images were captured using an Olympus SZX12 stereomicroscope, equipped with the Olympus MicroFire digital microscope camera U-CMAD3. Epifluorescence images were captured using an upright Olympus BX51 microscope (Tokyo, Japan), mounted with an Olympus XM10 camera. For confocal imaging, the ZEISS LSM 880 and 710 systems were used. Images were processed using Fiji (NIH), Adobe Illustrator 2020 and Adobe Photoshop CS6 and 2020 software.

### Quantification analyses

For quantification of CCND1-labeled cells, TUNEL, PHH3-labeling, and EdU incorporation, labeled cells were counted in cryostat sections midway through the optic cup (see Suppl. Fig. 1A for sagittal section scheme). The optic cup was sub-divided into 6 regions of interest (see Suppl. Fig. 3C) allowing comparison of dorsal and ventral individual subdivisions (D1, D2, D3 and V1, V2, V3) as well as combined data subdivisions to generate values for dorsal (D1-D3) and ventral optic cup domains (V1-V3). The percentage of cells was calculated by determining the number of total cells using DAPI-labeled nuclei. Cell diameter measurements were performed on whole eye images (examples are shown in Fig. 1C, D) by drawing a horizonal line across the center of the eye globe, measured using Fiji software (NIH). We analyzed a minimum of 6 eyes per genotype from n = 6 embryos (5 independent litters). To calculate the extent of attachment between OF margins, sagittal cryostat sections labeled with Phalloidin were analyzed. Attachment was defined for tightly apposed OF margins enclosing an area with phalloidin labeling apparent mostly as a single border between the OF margins (for examples, see Fig. 3A; Fig. 6C, D; Suppl. Fig. 3F). No attachment (set as “0”) was defined as a distinguishable gap between the two opposing phalloidin-labeled OF margins, with detectable mesenchymal cells present in the OF (for example, see Fig. 3B). Attachment of OF margins was measured by drawing a straight vertical line, analyzed using Fiji software. We analyzed 9 eyes; n = 7 embryos from 3 independent litters. Cellular height was calculated using Phalloidin-labeled images and Fiji software. Specifically, a straight line was applied along the apicobasal extent of each cell (Suppl. Fig. 3F), and 2-4 cells for each temporal or nasal margin of the OF was measured for each sample. At least 6 eyes from 5 embryos per genotype, from a minimum of 3 independent litters were analyzed. Statistical analysis was performed using Prism version 8 (Graphpad, San Diego, CA) for 2-Way ANOVA with Sidak’s multiple comparison or Student’s T-test. Differences were only considered significant when p < 0.05.

To quantify nuclear fluorescence intensity, YAP1 immunofluorescence confocal images of E11.5 cryosections midway through the optic cup were taken at 63x magnification. Two images were taken on either side of the optic fissure in the ventral region of the eye for both *Nf2*^CKO^ and control tissue. At least 42 nuclei in the RPE and at least 194 nuclei in RPE + retina combined were analyzed per eye. Three control eyes and four *Nf2^CKO^* eyes from different embryos were quantified. FIJI software was used to quantify mean nuclear YAP intensity. For quantification, we selected one slice of the z-stack (0.5 μm thickness) that was in the center of the nuclear plane of the optic cup. Background was subtracted using the FIJI rolling ball method. Nuclei in the ventral RPE and retina were manually traced using DAPI-counterstained images and added to the ROI manager. The nuclear ROI traces were then overlaid on their respective YAP1 channel to measure the area, mean intensity, and integrated density of the ROIs. For statistical analysis, average ROI (nuclei) area and average YAP1 integrated density per nuclei were used and unpaired two-tailed Welch’s t-tests were performed.

## Acknowledgements

The authors thank Katrina Hofstetter and Burns Newsome for technical support and Marco Giovannini, Ed Levine and members of the Levine and Fuhrmann laboratories as well as Jenny C. Schafer for helpful comments. We thank Jin Woo Kim for sharing data, Edward Levine for providing the Mitf^Mi^ allele and Marco Giovannini for providing the conditional Nf2 allele. This work was supported by the National Institutes of Health (R01 EY024373 to S.F, T32 HD007502 Training Grant in Stem Cell and Regenerative Developmental Biology to S.R, Core Grant P30 EY008126); a Catalyst Award to S.F. from Research to Prevent Blindness Inc./American Macular Degeneration Foundation, an unrestricted award to the Department of Ophthalmology and Visual Sciences from Research to Prevent Blindness, Inc.; the Vanderbilt University Medical Center Cell Imaging Shared Resource Core Facility (Clinical and Translational Science Award Grant UL1 RR024975 from National Center for Research Resources).

---

Supplemental Figure 1

**A:** Cartoon of scheme for sagittal sections. B, C: Low magnification images of E13.5 bright field of control eye (B) and *Nf2^CKO^* optic cup (C). In *Nf2^CKO^* optic cups, pigmentation extends deep into the unfused OF (C; arrowhead) and the RPE appears darker (C; arrow; compare with B). D, G: WGA labeling of the cell membrane demonstrates that cells in the E11.5 optic cup have little cytoplasm (Blue: Dapi). E, H: DAPI-stained nuclei were outlined in the RPE (yellow outlines) and the retina (not shown). F, I: Nuclei outlines (yellow) from E, H were overlayed on YAP1-stained images as ROIs and used for quantification. J: Average RPE nuclei (ROI) area demonstrate no difference between control and *Nf2^CKO^* nuclei. K: Average integrated density quantification in the RPE and retina combined displays significantly increased nuclear localization in *Nf2^CKO^* compared to control. L: Average nuclei area in the RPE and retina combined demonstrates no difference between control and *Nf2^CKO^* nuclei. Data are shown as means ± s.d., and p values are indicated above the horizontal lines in each graph (significant < 0.05). Statistical analysis was performed using unpaired, two-tailed Student T-test (Welch’s correction included for J-L). M, N: Lower magnification of YAP1 immunolabeling in E11.5 control and *Nf2^CKO^* embryos. YAP1 labeling in the lens and retina of controls (M; arrowheads). In *Nf2^CKO^*, YAP1 is detectable in nuclei of cells in the lens, retina, and RPE (N; arrowheads). Asterisks in D-I, M, N indicate OF.

Scale bars: 100 μm in B, C; 20 μm in D-I, 50 μm in M, N.

Supplemental Figure 2

Investigation of MITF protein localization and Mitf function in *Nf2^CKO^* eyes. A, B: At E11.0, MITF expression appears similar in the closing OF in control (A) and *Nf2^CKO^* eyes (B; arrowheads). Arrow in B points to MITF expression in the periphery. C-H: Mitf haploinsufficiency does not rescue coloboma in the *Nf2^CKO^* optic cup. C, D: In control eyes, removal of one Mitf allele results in reduction of pigmentation (D). G: Sagittal view of *Nf2^CKO^* optic cup showing persistent laminin expression ventrally (also shown in Fig. 1F at higher magnification). In *Nf2^CKO^;Mitf^Mi+/-^* optic cups, OF closure is not prevented, confirmed by persistence of laminin expression in the ventral optic cup (H). I, J: Analysis of LEF1 protein expression shows no change in the mutant RPE in the ventral optic cup (J). Arrowheads (E, F) and asterisks (A, B, G, H, I, J) label the OF. Sagittal view with nasal orientation to the left.

Scale bars: 50 μm in A, I, 200 μm in C, G.

Supplemental Figure 3

Analysis of apoptosis, eye diameter and height measurements in the RPE in *Nf2^CKO^* optic cups. A-D: TUNEL-labeled cells in the OF in E11.5 controls (A; arrows) and in *Nf2^CKO^* (B; arrows). Asterisks mark the overlying optic fissure. C: Scheme of subdivisions used for quantification in the optic cup. D: Quantification of TUNEL-labeled cells shows no significant change in *Nf2^CKO^* optic cups. E: The total number in ventral regions was quantified at E11.0 using Dapi labeling (see scheme in C). V1+3 contains counts of the two ventrolateral compartments and shows a significant increase in total cell number in *Nf2^CKO^*. V1-3 shows cell number counts for all three ventral regions combined. The region harboring the OF (V2) can exhibit a clear gap in *Nf2^CKO^* optic cups, therefore, the cell number can be underestimated. However, the total cell number shows still a significant increase, when V2 is included. F: No significant difference of the eye diameter is detectable between controls and *Nf2^CKO^* embryos. G: Example for measuring the apicobasal height of RPE cells in the nasal and temporal OF margins during alignment (E11.5). Measurements were performed using ImageJ. Asterisks mark the overlying optic fissure. H: Nested T-test analysis reveals no significant change in cellular height along in the apicobasal axis in cells lining the nasal OF margins in *Nf2^CKO^* embryos. Data are means (with minimal and maximal points). Statistical analysis for data shown in D and F was performed using Student T-test and in E using 2-Way ANOVA with Sidak’s multiple comparison. Data are means ± s.d., and p values are indicated above the horizontal lines in each graph. Sagittal view with nasal orientation to the left.

Scale bars: 100 μm in B, C.

## Abbreviations

AJ: Adherens junctions
Axin2: Axis inhibition protein 2
BHLH: Basic helix-loop-helix
BrdU: Bromodeoxyuridine
CCND1: Cyclin D1
Crb1: Crumbs family member 1
DAPI: 4’,6-diamidino-2-phenylindole
DAPL1: Death associated protein-like 1
DCAF1: DDB1 and CUL4 associated factor 1
ECM: Extracellular matrix
ERM: Ezrin, radixin, and moesin
FAT1: FAT atypical cadherin 1
FERM: Ezrin, radixin, and moesin
FGF: Fibroblast growth factor
LATS1/2: Large tumor suppressor 1/2
LEF1: Lymphoid enhancer binding factor 1
LRP6: Low density lipoprotein receptor-related protein 6
MAC: Microphthalmia, anophthalmia and coloboma
MITF: Microphthalmia associated transcription factor
MST1/2: Macrophage stimulating 1/2
NF2 (disorder): Neurofibromatosis Type 2
NF2 (gene): Neurofibromin 2
OF: Optic fissure
Otx2: Orthodenticle homeobox 2
PAX2: Paired box 2
PAX6: Paired box 6
PFA: Paraformaldehyde
PHH3: Phospho-histone H3
pMLC2: Phospho-myosin light chain II
PORCN: Porcupine
Rd8: Retinal degeneration 8
RPE: Retinal pigmented epithelium
SCRA: Sveinsson’s chorioretinal atrophy
TAZ: Tafazzin
TBX5: T-box 5
TEAD: TEA domain family member
TFAP2A: Transcription factor AP-2 Alpha
TRITC: Tetramethylrhodamine
TUNEL: Terminal deoxynucleotidyl transferase dUTP nick end labeling
VSX2: Visual system homeobox 2
WGA: Wheat Germ Agglutinin
Wnt: Wingless-type
YAP: Yes-associated protein
ZO-1: Zonula occludens-1

