## Supplementary figures and images for "Nf2 fine-tunes proliferation and tissue alignment during closure of the optic fissure in the embryonic mouse eye"

### Suppl. Fig.1

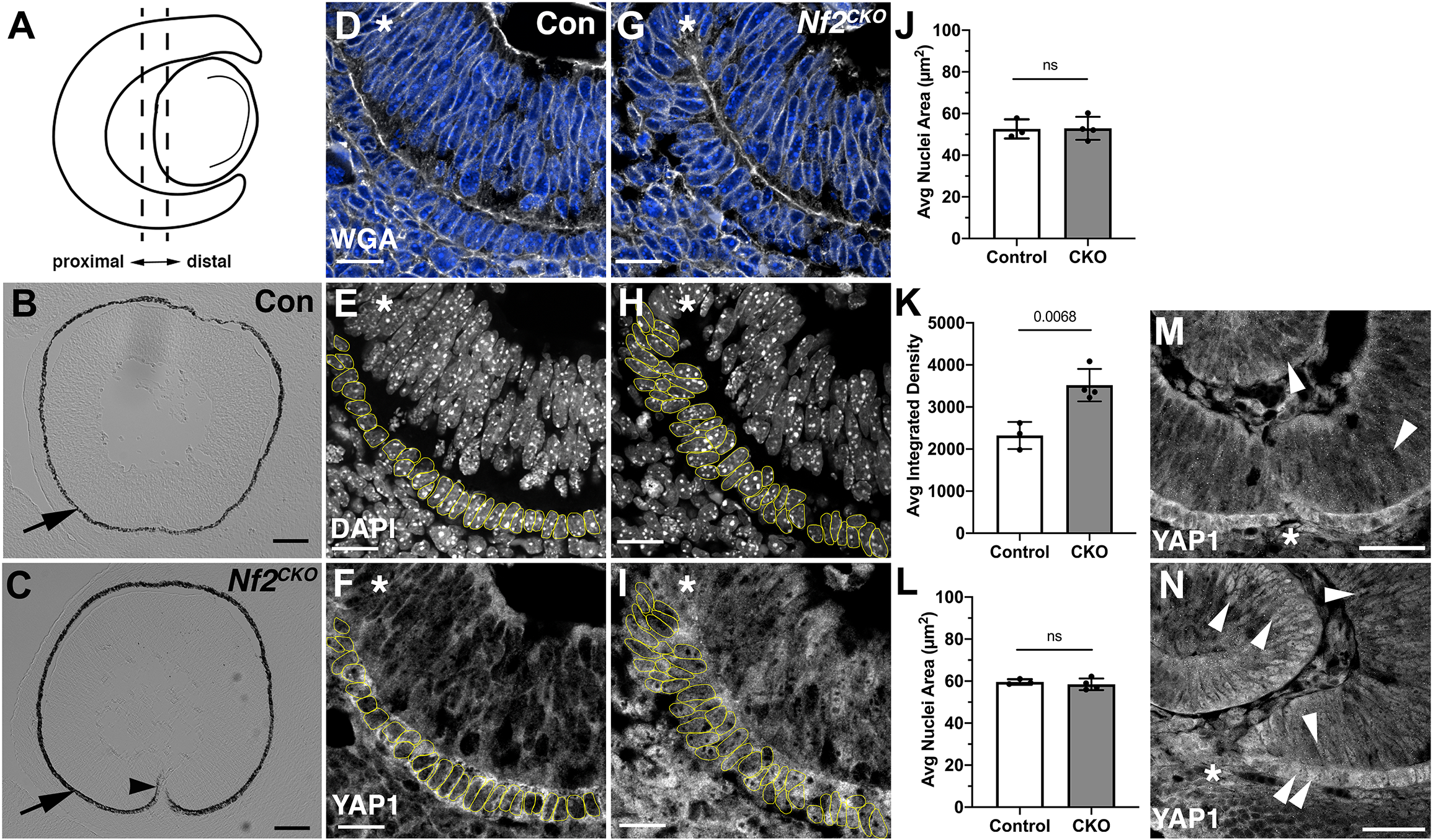

### Suppl. Fig.2

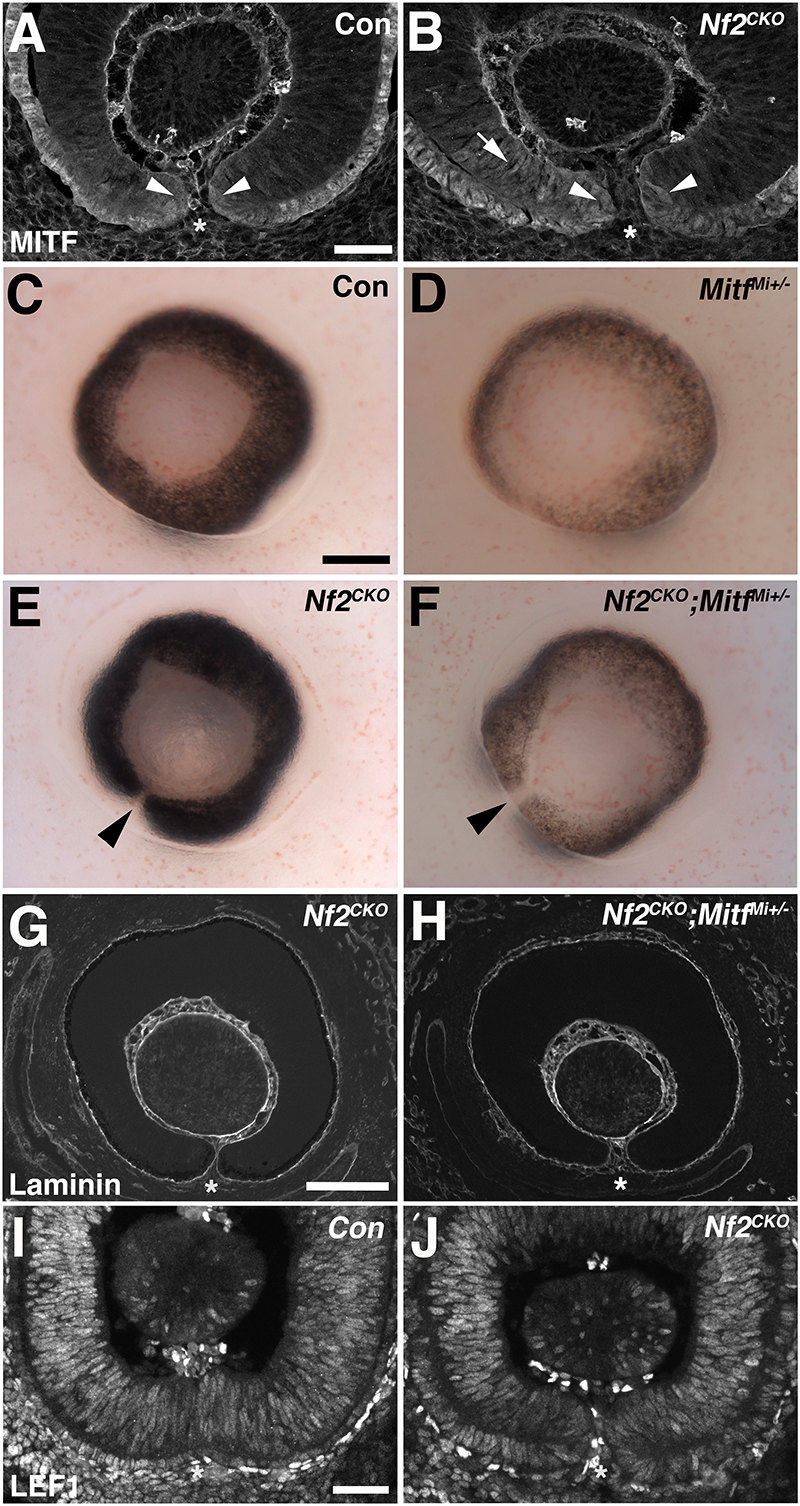

### Suppl. Fig.3

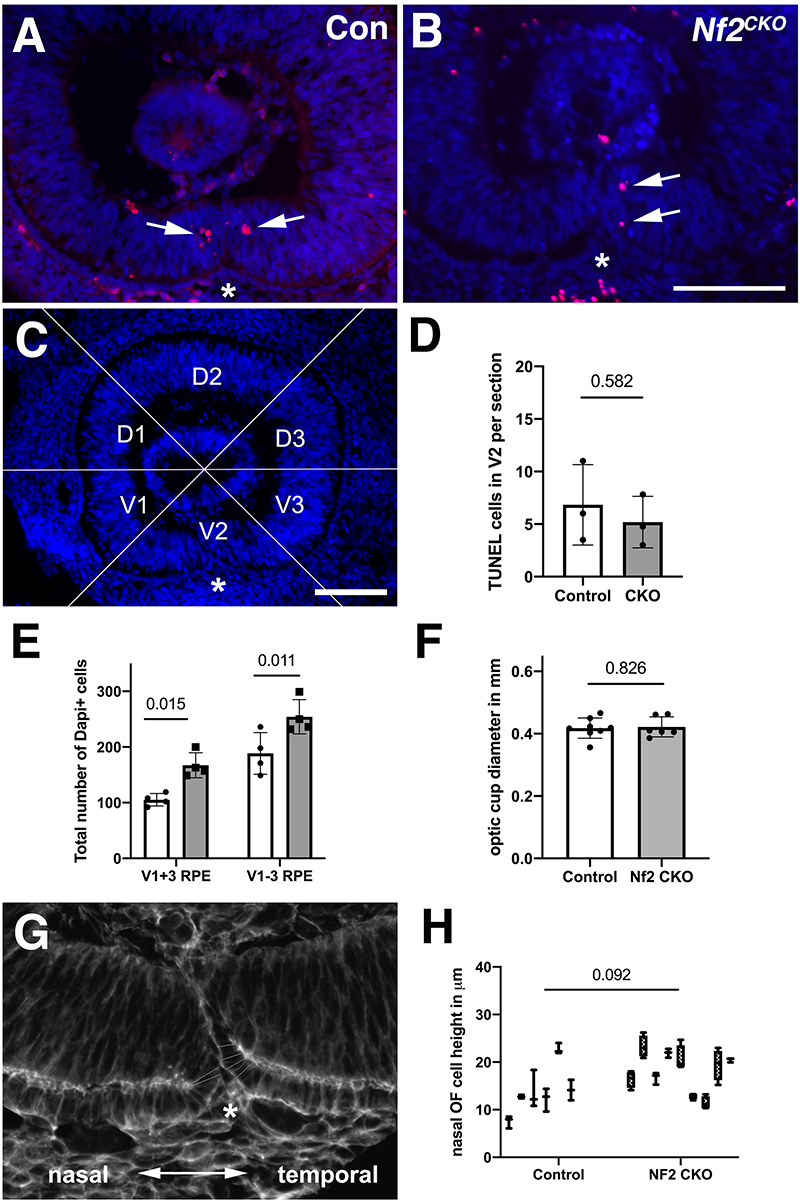
